# A potential cost of evolving epibatidine resistance in poison frogs

**DOI:** 10.1101/2023.01.04.522789

**Authors:** Julia M. York, Cecilia M. Borghese, Andrew A. George, David C. Cannatella, Harold H. Zakon

## Abstract

**Background:** Some poison arrow frogs sequester the toxin epibatidine as a defense against predators. We previously identified a single amino acid substitution (S108C) at a highly conserved site in a neuronal nicotinic acetylcholine receptor (nAChR) ß2 subunit that prevents epibatidine from binding to this receptor. When placed in a homologous mammalian nAChR this substitution minimized epibatidine binding but also perturbed acetylcholine binding, a clear cost. However, in the nAChRs of poison arrow frogs, this substitution appeared to have no detrimental effect on acetylcholine binding and, thus, appeared cost-free.

**Results:** The introduction of S108C into the α4β2 nAChRs of non-dendrobatid frogs also does not affect ACh sensitivity, when these receptors are expressed in *Xenopus laevis* oocytes. However, α4β2 nAChRs with C108 had a decreased magnitude of neurotransmitter-induced currents in all species tested (*Epipedobates anthonyi*, non-dendrobatid frogs, as well as human), compared with α4β2 nAChRs with the conserved S108. Immunolabeling of frog or human α4β2 nAChRs in the plasma membrane using radiolabeled antibody against the β2 nAChR subunit shows that C108 significantly decreased the number of cell-surface α4β2 nAChRs, compared with S108.

**Conclusions:** While S108C protects these species against sequestered epibatidine, it incurs a potential physiological cost of disrupted α4β2 nAChR function. These results may explain the high conservation of a serine at this site in vertebrates, as well as provide an example of a tradeoff between beneficial and deleterious effects of an evolutionary change. They also provide important clues for future work on assembly and trafficking of this important neurotransmitter receptor.

## Background

Some animals sequester alkaloid toxins for defense. Many of these alkaloids target ion channels, ion pumps, or neurotransmitter receptors (1, 2). Because the sequestered alkaloids are present within their tissues, defended animals (or their predators) must evolve protection from these toxins. This frequently occurs by adaptive amino acid substitutions at the target molecule (1–3), which may come with a cost of decreased function of that molecule (4–6) and decreased organismal performance. Because there may be a tradeoff between successful defense against predators and fitness (7), an important step in understanding the adaptive value of a sequestered toxin is a determination of the proximate cost of evolving target resistance.

Numerous neotropical poison frog species derive alkaloids from their arthropod prey, typically ants and mites, and sequester these toxins for defense (8–10). For example, at least three genera sequester alkaloid agonists or antagonists for nicotinic acetylcholine receptors (nAChRs) which, in turn, disrupts acetylcholine-based synaptic transmission. Frogs, like mammals, have numerous cholinergic neurons in their brains (11, 12) and some species have evolved resistance to their dietary alkaloids. In particular, species of *Epipedobates* poison frogs sequester the alkaloid epibatidine, well-studied for its agonist action on nAChRs that contain β2 subunits, and we previously demonstrated epibatidine-resistance in at least one brain-expressed isoform of the nAChR (13).

nAChRs exist as a diverse family of molecules composed of different pentameric combinations of homologous subunits derived from at least 17 genes (α1-α10, β1-β4, γ, δ, ε). The properties of nAChRs are determined by their subunit composition, giving rise to multiple subtypes with a range of overlapping pharmacological and biophysical properties (14). In mammals, the major brain isoform of the nAChR is composed of α4 and β2 subunits (Additional file 1). The most common stoichiometry is two α4 and three β2 subunits (2α:3β), which produces an isoform of the receptor with high sensitivity (HS) to ACh and where ACh binds at α(+):β(-) interfaces (Additional file 1, panel A). However, a low-sensitivity (LS) binding site is also naturally present (15) when the receptors within the central nervous system express three α4 and two β2 subunits (3α:2β, Additional file 1, panel B). LS sites, which occur at the interface between two adjacent α4 subunits [i.e. α(+):α(-)] (16, 17), can influence the function of the pentameric α4β2 nAChR by providing an additional low-affinity binding site for ACh (18, 19).

Experimental expression of nAChRs in *Xenopus* oocytes has been critical for understanding their normal function (15, 20), pathology (21–23), and pharmacology, including responses to epibatidine (24, 25), because many of their properties (as measured *in vivo*) can be replicated in this heterologous system. Pertinent to this study, the naturally occurring differences in ACh sensitivity can be replicated in *Xenopus* oocytes by varying the RNA ratio of α4 and β2 subunits (26, 27). An abundance of α4 RNA favors 3α:2β with both HS and LS binding sites, whereas an abundance of β2 RNA favors 2α:3β with only HS binding sites (Additional file 2, panels A and B). This observation has been verified with concatemers with fixed ratios of α4 and β2 subunits (28).

In our previous work, we noted a serine to cysteine substitution (S108C) that evolved independently in the β2 nAChR subunit in three genera of dendrobatid frogs (*Epipedobates, Ameerega, Oophaga*) that possess alkaloids which target α4β2 nAChRs (13) (Fig. 1). The serine at this site is highly conserved across vertebrate β2 nAChR subunits over 550 million years of evolutionary time (Fig. 1) (29). We found that introduction of the S108C substitution into the human α4β2 nAChR (1α:3β RNA ratio expressed in *Xenopus* oocytes, which should produce only HS sites) conferred epibatidine resistance (13) but also produced LS sites, reducing the sensitivity to ACh as detected in concentration response curves (CRCs) (Additional file 2, panel A). At a 3α:1β RNA ratio, the introduction of S108C added even more LS sites, further reducing the sensitivity to ACh by shifting the ACh CRC rightward (Additional file 2, panel B). We also noted substitutions at other highly conserved sites near S108C in other dendrobatids (*Epipedobates* and *Ameerega*) (Fig. 1). Introduction of F106L, a novel phenylalanine to leucine substitution in *Epipedobates*, into the human β2 nAChR subunit with the S108C substitution partially (3:1) or completely (1:3) restored baseline ACh sensitivity (i.e. LS sites were eliminated) (Additional file 2, panels A and B). Therefore, the novel presence of both L and C decreases sensitivity to epibatidine in human α4β2 nAChRs while maintaining a normal response to ACh, which would presumably be advantageous for frogs defended by epibatidine.

**Fig. 1.**
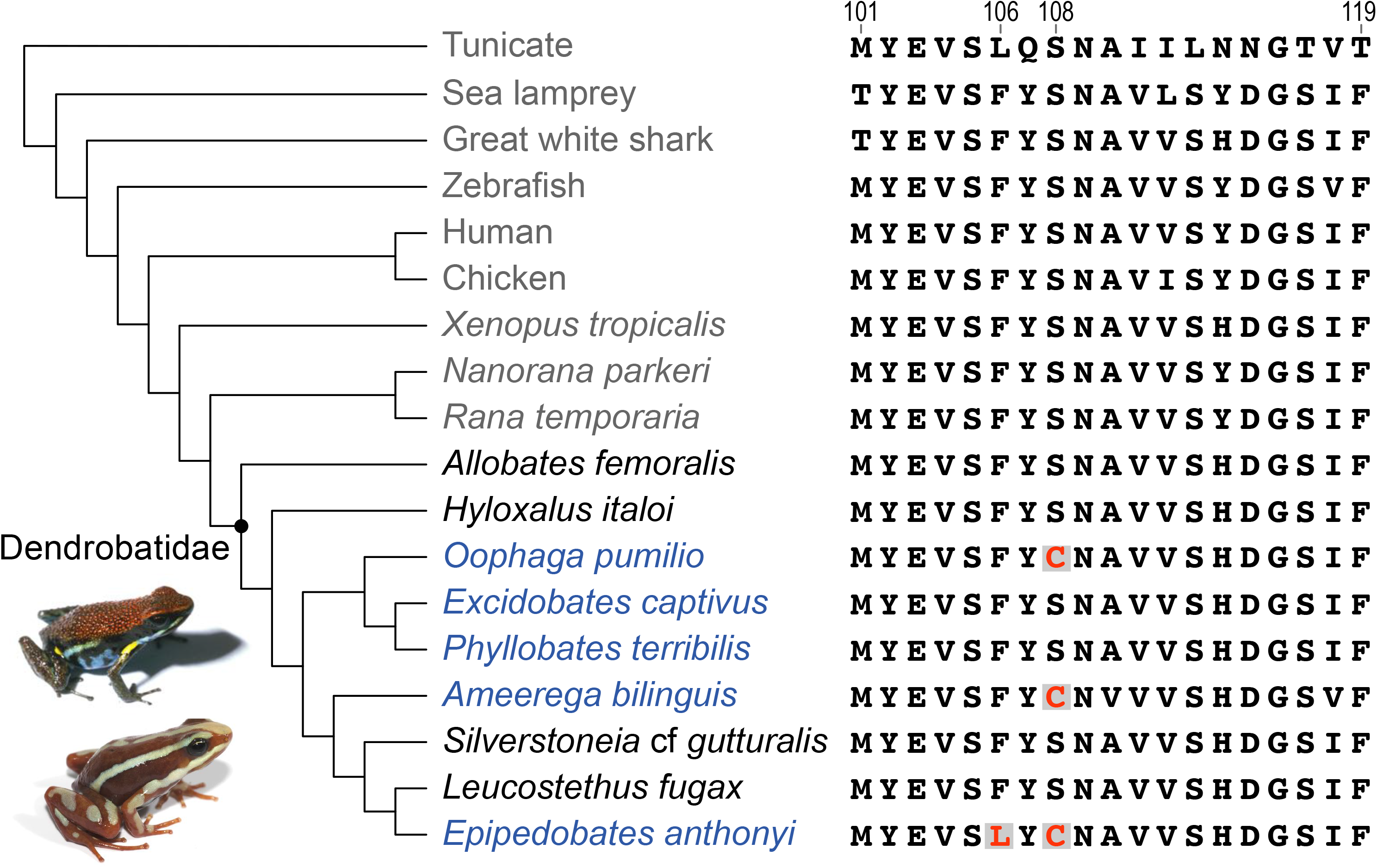
Phylogeny of selected chordates showing the variation of amino acid sequences in the region of interest of the β2 nAChR subunit. Scientific names are used for the frogs. The dot in the phylogeny represents the ancestor of the poison frogs (Dendrobatidae clade). The names of undefended species of poison frogs are in black and those of defended species are in blue. Defense by sequestered alkaloids has evolved three times, associated with parallel evolution of S108C and the unique evolution of F106L in *Epipedobates*. Photos of *Ameerega bilinguis* and *Epipedobates anthonyi* (from which epibatidine was first isolated) are shown in the lower left.

Having tested the response in human α4β2 nAChRs, we predicted that the wild-type *Epipedobates* receptor, which evolved the S108C and F106L substitutions, would likewise possess resistance to epibatidine and incur a similar decrease in ACh sensitivity (due to the C108), but this decreased ACh sensitivity would be rescued by its second substitution (leucine instead of the ancestral phenylalanine) (13). However, while the *Epipedobates* wild-type α4β2 nAChR (expressed after injection of either 1:7 or 7:1 α4:β2 RNA ratio) showed epibatidine resistance (13), there was surprisingly little decrease in the ACh sensitivity between the two ratios (Additional file 2, panels D and E), even when RNA was injected in a ratio that strongly favors the formation of LS sites in mammalian nAChRs (7α:1β) (Additional file 2, panel E). Reverting C108 to the ancestral serine (S) eliminated epibatidine resistance but did not decrease ACh sensitivity (in either 1:7 or 7:1 α4:β2 RNA ratios). Additionally, reverting the leucine at position 106 to the ancestral phenylalanine had no effect either on the ACh (Additional file 2, panels D and E) or the epibatidine CRCs observed in *Epipedobates* α4β2 nAChRs (13). This evidence contradicted our initial hypothesis that the cost of epibatidine resistance for poison frogs is decreased ACh sensitivity.

Here, we test three alternative hypotheses for the unexpected inability of the C108S substitution to alter the ACh sensitivity of the *Epipedobates* nAChR: 1) other unidentified dendrobatid-specific amino acid substitutions protect the receptor; 2) the cost manifests as a different aspect of receptor function or 3) this substitution is cost-free. In comparing α4β2 nAChRs from *Epipedobates*, non-dendrobatid frogs, and humans, we found no support for additional dendrobatid-specific substitutions (rejection of hypothesis 1) but observed instead that S108C presents a differential potential cost (rejection of hypothesis 3): it decreases the number of α4β2-containing nAChRs in the plasma membrane (failure to reject hypothesis 2), which could potentially disrupt cholinergic synaptic transmission, presumably leading to a decrease in fitness.

## Results

### The S108C Substitution Does Not Affect Acetylcholine Sensitivity in Non-Dendrobatid Frogs

We tested the “unidentified dendrobatid-specific substitution” hypothesis by examining the effect of the S108C substitution on the ACh sensitivity of the α4β2 nAChR of two deeply divergent (30) species of non-dendrobatid frogs –Western clawed frogs (*Xenopus tropicalis*, 182 million years ago, mya) and high Himalaya frogs (*Nanorana parkeri*, 130 mya)—with the expectation that their receptors would behave more like human receptors than those of dendrobatids.

The ACh CRC from *Xenopus* α4β2 nAChRs was best fit by a monophasic curve with a single EC_50_ (concentration of transmitter that elicits 50% of the maximum response), whether ratios favored (1:3) or disfavored (7:1) the incorporation of β2 subunits. The monophasic curve implies a single population of HS receptors with only HS binding sites (Fig. 2A-C; Additional files 6 and 9). LS sites were never observed with either ratio, unlike homologous mammalian receptors that we previously tested (Additional file 2, panel B). We did not detect differences in the ACh CRCs between the *Xenopus* wild-type [with phenylalanine in position 106 and serine in position 108 in the β2 subunit, represented with β2(FS)], the S108C-substituted [β2(F**C**), with bold indicating the substituted residue], or the combined F106L/S108C-substituted nAChRs [β2(**LC**)]. Interestingly, the *Xenopus* α4β2 nAChR was ~28x more sensitive to ACh than the human receptors (Additional files 6 and 8, and (13)), and ~ 9x more sensitive than the *Epipedobates* receptors (13). This was true even when human and *Xenopus* receptors were tested in the same recording session with the same reagents.

**Fig. 2.**
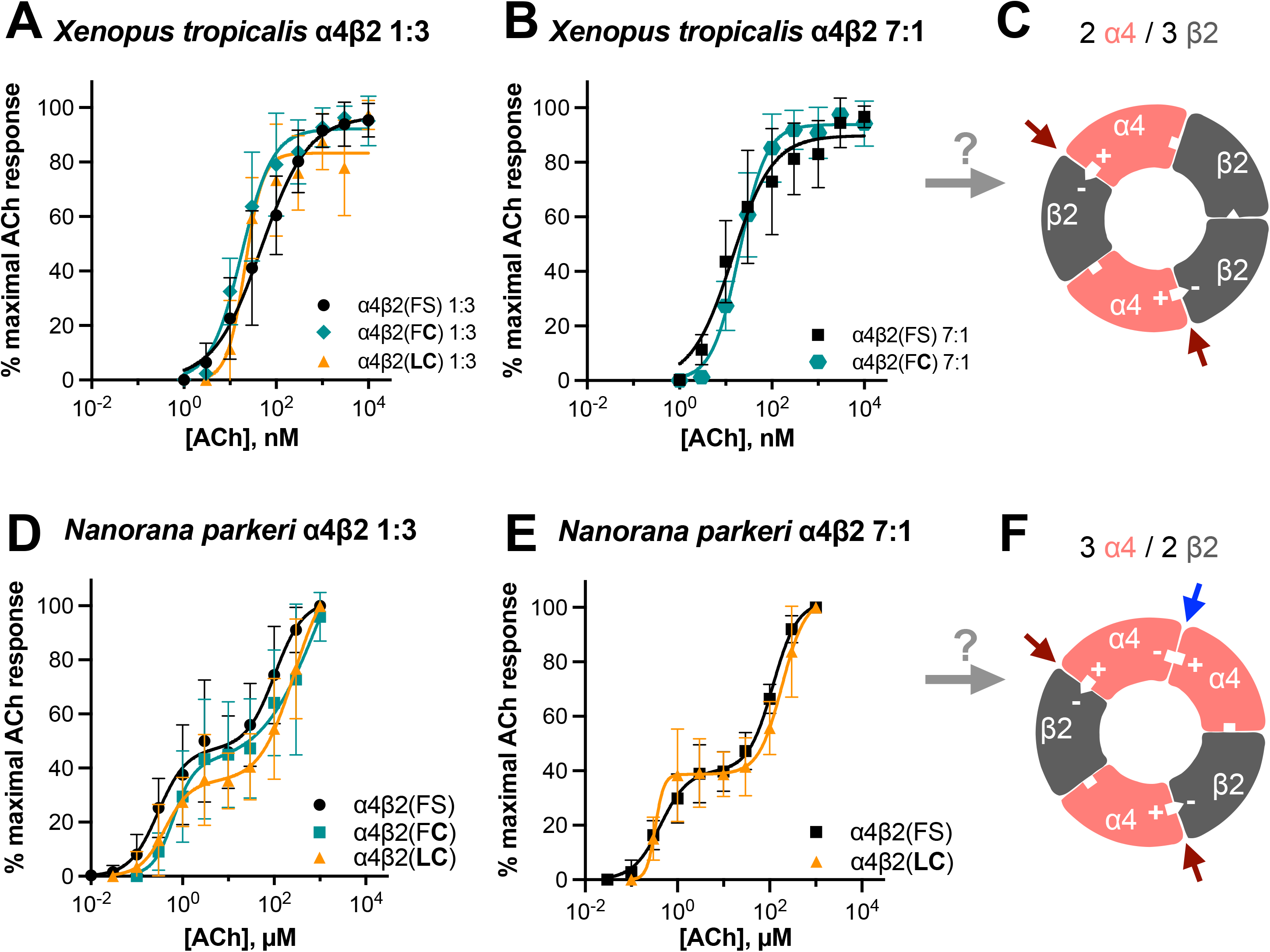
Acetylcholine CRCs of receptors from two non-dendrobatid frogs. *Xenopus tropicalis* retain a monophasic ACh CRC best fit with a single EC_50_ (Additional files 6 and 9) even in conditions that induce LS sites in mammalian α4β2 nAChRs (7α:1β) or with the S108C substitution alone [β2(F**C**)] or in combination with F106L [β2(**LC**)]. The α:β RNA ratios are 1:3 (A, n= 6-18) and 7:1 (B, n= 13-37). The actual stoichiometry of frog receptors is unknown but the *Xenopus* α4β2 nAChR behaves as the mammalian 2α:3β stoichiometry (C). This conjecture is indicated as a question mark over the grey arrow. Note that the scale for *Xenopus* is nanomolar concentration. *Nanorana parkeri* retain a biphasic CRC best fit with two EC_50_ values (Additional files 7 and 9) in both 1:3 (D, n= 5-9) or 7:1 (E, n= 6-9) α:β RNA ratios, with the S108C substitution alone [β2(F**C**)] or in combination with F106L [β2(**LC**)]. The *Nanorana* α4β2 nAChR behaves as if its stoichiometry is 3α:2β (F) with both ratios of RNA used in this study. Data points represent means ± SD. Red arrows indicate HS binding sites and blue arrow indicates LS binding site. **+** and – signs indicate the principal and complementary components of the subunit interfaces.

In contrast, the ACh CRC from *Nanorana* α4β2 nAChRs was biphasic and best fit with two EC_50_ values (Fig. 2D and E; Additional files 7 and 9), implying the presence of both LS and HS α4β2 isoforms (Fig. 2F). When recording the aggregate responses of many receptors, it is not possible to distinguish between all receptors having both HS and LS binding sites versus two populations of receptors, one with HS sites and another with LS sites. However, these alternatives may be disambiguated by varying the α:β ratios. We observed that the ACh CRCs were biphasic in all α:β ratios, implying that a single *Nanorana* α4β2 nAChR possess both HS [α(+):β(-)] and LS [α(+):α(-)] sites.

These data demonstrate that the cysteine substitution (S108C) alone or in combination with F106L does not affect ACh sensitivity in non-dendrobatid frogs, leading us to reject the hypothesis that dendrobatid-specific substitutions in the α4β2 nAChR protect against perturbation to CRCs by preventing the ratio-dependent emergence of LS sites as occurs in mammalian receptors. Further, they suggest that, unlike mammalian α4β2 nAChRs, the frog α4β2 nAChRs that we studied have fixed stoichiometries of α and β subunits that cannot be altered by skewing the ratio of their RNAs, at least in the oocyte expression system and with the ratios of RNAs that we used (Fig. 2C and F). Finally, they show that frogs have surprising but as yet unexplained species-specific diversity in the ACh CRCs of their α4β2 nAChRs.

### S108C Substitution Reduces Current Magnitudes of Expressed Frog α4β2 nAChRs

Our second hypothesis is that S108C might perturb a different aspect of receptor function than ACh sensitivity. While measuring CRCs, we noted that currents from *Xenopus* or *Nanorana* α4β2 nAChRs with the S108C substitution, especially in the 7α:1β ratio, were absent or too miniscule to reliably measure using amounts of RNA that generated substantial currents in wild-type channels. Therefore, especially at ratios of 7α:1β, we increased the total amount of RNA to enhance current amplitudes (Additional file 3). The observation that cysteine-bearing receptors had smaller currents than wild-type receptors, even when higher amounts of RNA were injected, suggested that the presence of cysteine reduces the amplitude of α4β2 nAChR currents.

For example, in *Nanorana*, at 1:3 or 7:1 α4:β2 RNA ratios, S108C with or without F106L reduced current amplitudes in both ratios (Additional file 3, panel B). Similar to *Nanorana*, we observed in the *Epipedobates* receptor that S108C with or without F106L had a significantly reduced maximal current (Fig. 3A). The observations in *Epipedobates* are particularly revealing as they show that adding the ancestral and conserved amino acid, serine, increases receptor currents compared with the *Epipedobates* wild-type with C108. Thus, a major effect of the S108C substitution in the β2 nAChR subunit is to decrease maximal currents compared with serine-containing β2 subunits obtained with similar amounts of RNA. Such a decrease in macroscopic currents would potentially compromise synaptic transmission, with presumed downstream effects on organismal function.

**Fig. 3.**
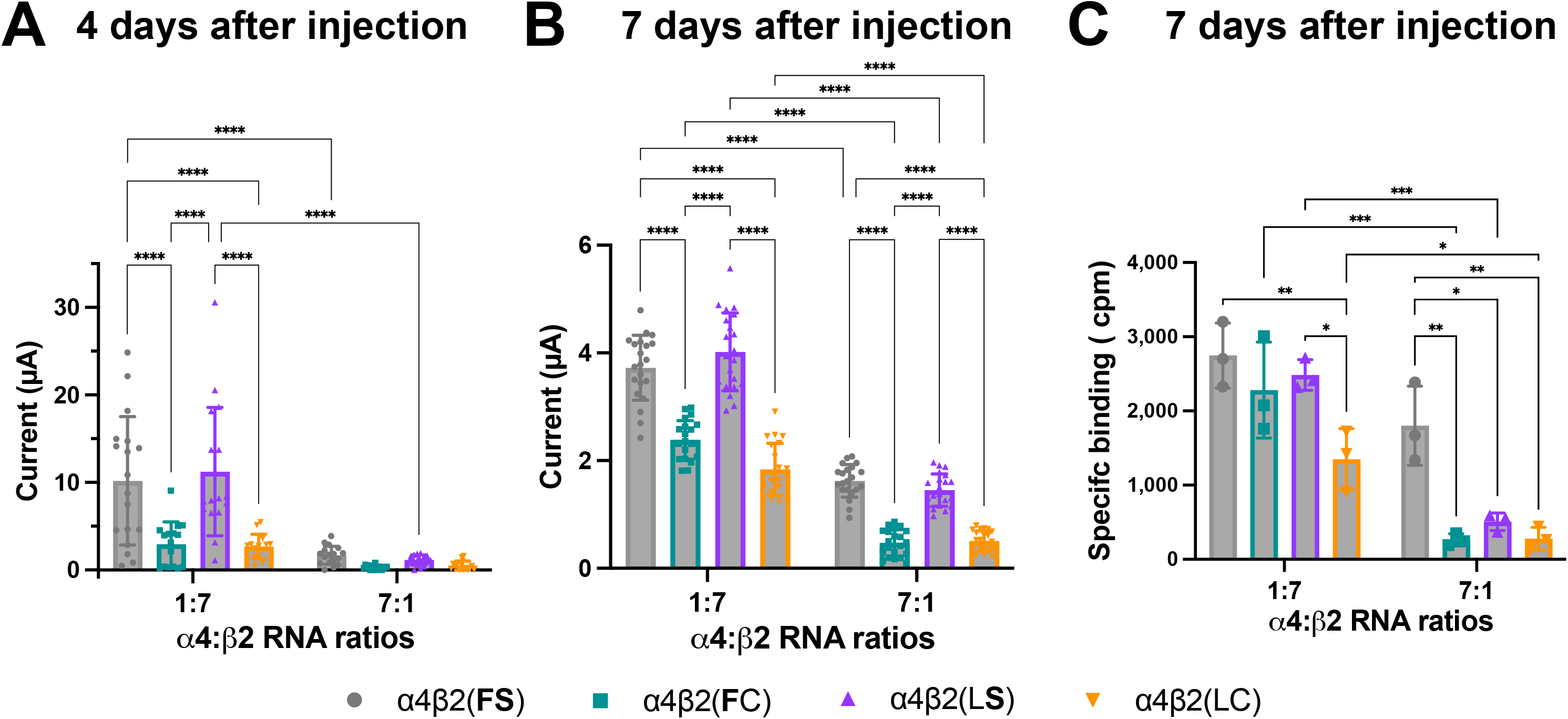
Maximal ACh-induced current and labeling of *Epipedobates anthonyi* α4β2 nAChRs. The presence of cysteine in location 108 decreases the magnitude of ACh-stimulated maximal currents, either with F106 [β2(**F**C)] or with L106 [β2(LC), wild type] (A, n= 14-18 oocytes; B, n= 21). The bold font indicates a substitution introduced in the wild type receptor. Following recordings shown in B, oocytes were treated with the radiolabeled antibody ^125^I-mAb 295 to quantify the specific binding, i.e. the number of β2-containing nAChRs expressed in the plasma membrane (C, n= 3, 7 pooled oocytes each experiment). Cpm stands for counts per minute. α4:β2 RNA injection ratios favoring HS (1:7) and LS (7:1) stoichiometries are indicated on the X-axis. Data is presented as means ± SD, and were analyzed using two-way ANOVA, followed by pairwise comparisons corrected for multiple comparisons with Holm-Šídák’s test. *\*P*< 0.05, \*\**P*< 0.01, \*\*\**P*< 0.001, \*\*\*\**P*< 0.0001.

### S108C Decreases the Number of nAChRs in the Plasma Membrane

In principle, a decrease in current magnitude induced by the cysteine substitution could be due to 1) a reduction in the total number of functional receptors on the cell surface or 2) changes in the biophysical/kinetic properties of the expressed receptors. We tested the first alternative with *Epipedobates* α4β2 nAChRs by measuring the number of receptors in the plasma membrane of intact oocytes labeled with a radiolabeled antibody (^125^I-mAb 295) that specifically binds to the β2 nAChR subunit of mature α4β2 nAChRs (31). Additional file 1, in panels C-E shows a fragment from monoclonal antibodies that similarly binds to the β2 subunit of human α4β2 nAChRs (32). The iodinated antibody binds to chicken and mammalian receptors (33), and we now extend the species range by showing that it binds to frog receptors as well (Additional file 4, panels A,B). Additionally, the strong relationship (R^2^=0.83) between maximum current and specific binding determined as described in the following paragraphs is further evidence that this antibody binds to frog nAChRs (Additional file 4, panel C).

We injected RNAs encoding single *Epipedobates* α4 and β2 nAChR subunits into oocytes and allowed them to incubate for 7 days (31, 34). We confirmed that oocytes incubated for 7 days generate the same relative maximal currents across the different constructs as those incubated for 4 days (Fig. 3A and B). In general, the presence of C108 in the β2 nAChR subunit resulted in decreased macroscopic currents compared with β2 S108-containing nAChRs, independently of the presence of F or L in position 106 in the β2 subunit. Additionally, when the β2 subunit was the limiting factor (7α:1β ratio), the current was decreased compared with the 1α:7β ratio.

In a ratio favoring β2 subunit incorporation (1α:7β), the number of receptors per oocyte measured using ^125^I-mAb 295 was similar except for the wild-type *Epipedobates* receptor [β2(LC)], which was significantly lower (Fig. 3C). However, when β subunit concentration was limiting (7α:1β), the presence of serine in position 108 in conjunction with phenylalanine in position 106 [β2(**FS)**, that is, the ancestral residues present in human, *Xenopus* and *Nanorana* β2 subunits] significantly increased the number of α4β2 nAChRs in the plasma membrane (Fig. 3B and C). This supports the hypothesis that the presence of C108 reduces the total number of α4β2 nAChRs in the plasma membrane. It also suggests that the presence of L106 together with C108 is detrimental to receptor availability, which was unexpected since F106L was previously shown to compensate for the alterations in the ACh CRC caused by S108C (FS = **LC ≠** F**C**) in mammalian receptors (13).

### Concatenated Human nAChRs Confirm Results from Frogs

The most parsimonious explanation for our results with frog receptors is that the presence of 108C in β2 subunits limits the number of functional α4β2 nAChRs by, for example, decreasing the efficiency of receptor assembly, trafficking, or stability. Could such an effect also explain our previous data on human receptors? The S108C substitution in human β2 subunits biases the receptor toward more LS sites, especially when β2 subunits are scarce (7α:1β). The presence of 108C could either directly alter ACh sensitivity in a 2α:3β stoichiometry or, alternatively, induce the 3α:2β stoichiometry in which LS sites result from the inclusion of an extra α4 subunit in the face of less efficient β2 subunits. Injection of RNA into cells results in a mixed population of receptors with different stoichiometries. Biasing the ratios of RNA (e.g., 7:1) as we have done, strongly favors one stoichiometry over others. However, this is still not a pure population of HS or LS nAChRs. This can be overcome by using concatemeric constructs in which the number and order of α4 and β2 subunits are identical in all receptors because they genetically encoded, covalently-linked *α4* and β2 nAChR subunits (see methods). To test between these alternatives, we generated a series of concatenated human α4β2 nAChR constructs with fixed 2α:3β configurations, but with one, two, or all three β2 subunits containing the S108C substitution. If S108C induces LS sites in receptors composed of free subunits, with an LS composition (2α and 3β), the concatemeric receptors would produce a biphasic ACh CRC with no decrease of the current magnitude. If S108C produces a deficiency in the subunits’ ability to assemble into functional pentamers and/or traffic to the plasma membrane, then a monophasic ACh CRC (HS-like) and a decrease in current magnitude would be observed for the concatemeric receptors.

We found that the current amplitude of α4β2 nAChR was dramatically decreased with a single S108C-containing subunit, and no currents could be recorded when S108C was present in two or three subunits (Fig. 4A-C; Additional file 8). Importantly, the ACh CRCs generated from concatenated α4β2 nAChRs harboring a single S108C had no LS component (Fig. 4A). This supports the contention that the LS sites observed in human α4β2 nAChRs formed from free subunits (Additional file 2, panel A) derive from replacing a β2 subunit with an α4, rather than from the S108C substitution introducing LS sites in addition to the HS sites.

**Fig. 4.**
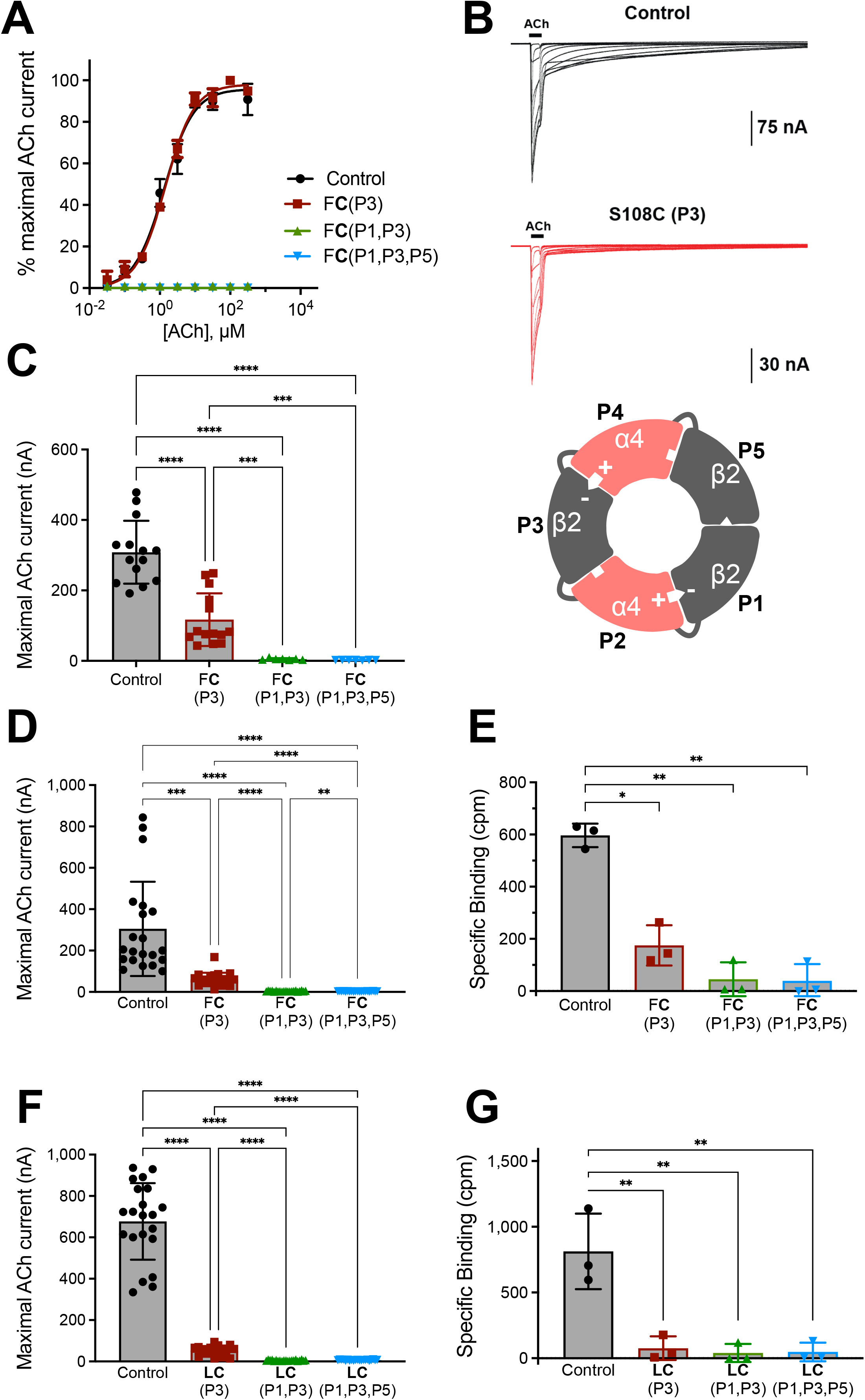
Currents and number of receptors measured after expression of concatemeric human nAChRs. All concatemers have an enforced 2α:3β stoichiometry (indicated by inset of schematic receptor). Wild-type receptor [F106 and S108, β2(FS)] generates a CRC with a monophasic fit (A, black). A similar concentration-response profile was observed for concatenated α4β2 nAChRs with a S108C substitution (A, red, F**C**, substitution indicated by bold font; P3 = position 3 within the linked receptor) in a single β2 subunit. However, concatenated α4β2 nAChRs with two (A, green) or all three (A, blue) β2 subunits containing S108C in the indicated positions generate no current. CRC analysis can be found in Additional files 8 and 9, *n=* 7-14. Raw current traces to increasing concentrations of ACh (B). Concatemers where a single β2 subunit has a S108C substitution [β2(F**C**)] show significantly reduced currents. As stated in panel A, concatemers with two or three S108C-containing β2 subunits generate no current (C, *n*= 7-14). Baseline recordings of maximal currents (D and F, *n*= 21) preceding harvesting of oocytes for measurements of receptor number (E and G, *n*= 3, 7 pooled oocytes in each experiment). Results for concatenated α4β2 nAChRs with a S108C substitution (F**C**) in D and E. Addition of F106L substitution to S108C [β2(**LC**)] did not rescue the effect of S108C (F, G). In all cases, concatenated α4β2 nAChRs with a single S108C-containing β2 subunit have significantly reduced numbers of receptors in the plasma membrane when compared to controls. Those concatenated α4β2 nAChRs with two or three β2 subunits harboring the S108C mutation are not expressed in the plasma membrane (i.e., the values were no different from un-injected oocytes). Specific binding was measured as counts per minute (cpm). Data are shown as means ± SD, and were analyzed using Brown-Forsythe and Welch ANOVA tests, followed by Dunnett’s T3 multiple comparisons test. * *P*< 0.05, \*\**P*< 0.01, \*\*\**P* < 0.001, \*\*\*\**P*< 0.0001.

Finally, in the same oocytes we measured both maximal ACh-induced currents and the number of α4β2 nAChRs in the plasma membrane. The maximal currents followed the same pattern previously found (Fig. 4D) and in agreement with the electrophysiology, that the total cell-surface expression fell with a single S108C-containing β2 subunit and were vanishingly small with two or three S108C β2 subunits incorporated into the functional pentamer (Fig. 4E). Also, in agreement with the data from frogs, the addition of the F106L substitution did not compensate for S108C, neither in the maximal currents (Fig. 4F) nor the total number of nAChRs present in the plasma membrane (Fig. 4G). The fact that S108C decreases current magnitude in frog and human concatenated α4β2 nAChRs emphasizes the general detrimental nature of this substitution on vertebrate α4β2 nAChRs.

## Discussion

One lineage of dendrobatid frogs, *Epipedobates*, sequesters the nAChR agonist epibatidine for defense (35). A β2 cysteine substitution that decreases the α4β2 nAChR sensitivity to epibatidine evolved three times in parallel in different groups of dendrobatids. Even though two of these three (*Oophaga* and *Ameerega*) are not known to sequester epibatidine, they sequester other nAChR agonists or antagonists. The serine at this site (108) is otherwise highly conserved in vertebrate β2 subunits, suggesting that substitutions at this site are maladaptive. Our previous work showed that, while the S108C substitution protected human receptors against epibatidine, it perturbed their ACh sensitivity, an obvious cost (Fig. 5). However, we also found that the S108C substitution had little to no effect on the ACh sensitivity of dendrobatid receptors, suggesting three possibilities: that this advantageous substitution is cost-free in frogs, that the cost manifests in other properties of nAChR function, or that there are additional, unrecognized dendrobatid-specific substitutions that prevent perturbation of ACh sensitivity (36).

**Fig. 5.**
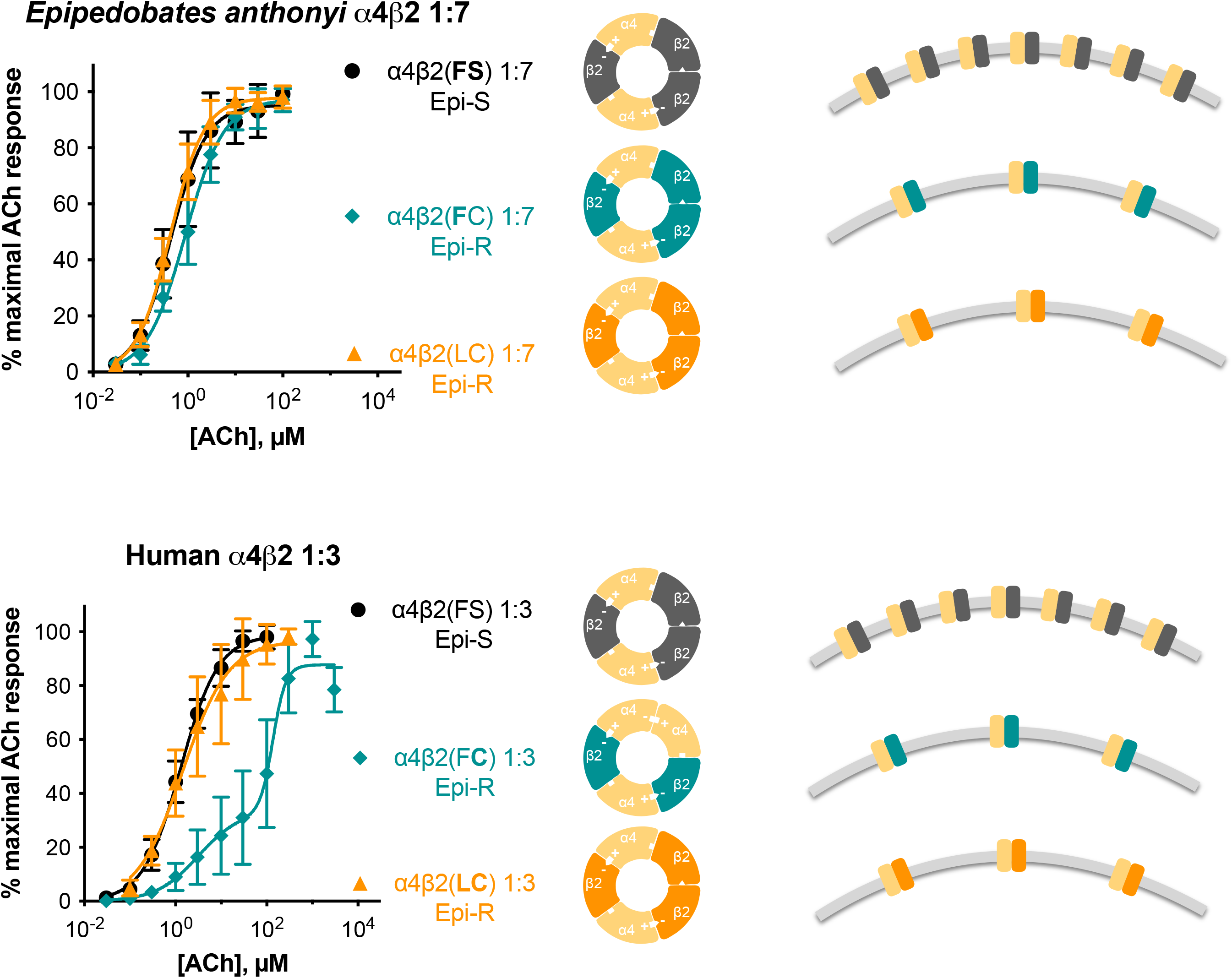
Summary figure. Results from Tarvin et al. (2017) are shown in the graphs on the left, from *Epipedobates anthonyi* (top) and human (bottom) nAChRs (RNA ratio 1α:7β for *Epipedobates* and 1α:3β for human). Epi-S and Epi-R refer to the epibatidine-sensitive and -resistant characteristics of the receptor. These results led us to hypothesize that the mutation S108C in the human β2 subunit [α4β2(F**C**)] resulted in an altered stoichiometry: instead of a monophasic curve (characteristic of α4β2 nAChRs composed of 2 α4 and 3 β2 subunits, the ACh concentration-response curve for this mutant was biphasic and shifted to the right (characteristic of α4β2 nAChRs composed of 3 α4 and 2 β2 subunits. Furthermore, this alteration in the ACh sensitivity was not observed in *Epipedobates* receptors, which showed the same monophasic curve at all RNA ratios tested (Additional file 2). In the central panels, the predicted stoichiometry is shown as a diagram of the receptor. We now report that there is a reduction of maximal ACh-induced currents in *Epipedobates* receptors with C108-containing β2 subunits, likely due to reduced availability of C108-containing β2 subunits (Fig. 3). The relative number of receptors in the plasma membrane is shown on the right diagrams. No differences in ACh sensitivity were observed after biasing α4 and β2 RNA injection ratios, indicating that *Epipedobates* α4β2 nAChRs functionally assemble in a single stoichiometry. However, the reduced cell-surface expression of α4β2(F**C**) nAChRs (also observed for *Epipedobates* α4β2 nAChRs containing cysteine in position 108 of the β2 subunit), alters the concentration-response profile of human α4β2(F**C**) receptors from the monophasic (HS-like) of the wild type nAChR, to biphasic (LS-like) CRC, indicating an alternative stoichiometry. The presence of an additional mutation [β2(**LC**)] confers an HS-like stoichiometry but does not correct the β2 reduced availability. The studies on the human receptor numbers in the plasma membrane were obtained using concatemeric receptors (Fig. 4).

We tested the “unknown dendrobatid-specific substitution” hypothesis by assessing concentration-response profiles of the endogenous neurotransmitter ACh of two phylogenetically divergent non-dendrobatid frogs, with the expectation that these species would show perturbations of ACh CRCs similar to those of human α4β2 nAChRs. But the ACh sensitivity of these frog nAChRs were also unaffected by the S108C substitution. Unexpectedly, however, in both dendrobatid and non-dendrobatid frogs, S108C induced a drastic reduction of ACh-activated maximal currents and the number of α4β2-containing nAChRs in the plasma membrane, exposing a potential detriment due to the S108C substitution. This is particularly notable in that introduction of the ancestral serine into *Epipedobates* β2 subunit markedly rescues current levels and numbers of α4β2 nAChRs in the plasma membrane over its wild-type cysteine-containing β2 subunit.

### The F106L Substitution Has Minimal Effect

Like S108, F106 is highly conserved in vertebrate β2 subunits (Fig. 1). We previously observed that F106L wholly or partially rescued the effects of S108C on the CRCs of human α4β2 nAChRs assembled from single nAChR subunits (36). In the present study we saw little effect, or occasionally even a detrimental effect, of F106L on current magnitude or receptor numbers with α4β2 nAChRs expressed from free frog or concatenated human subunits. One possibility is that F106L affects the biophysical properties of the receptors (e.g., single-channel open probability, open and/or closed dwell-times, etc.) which we did not study here, to counteract the detrimental effects of S108C. Additionally, *Epipedobates anthonyi* apparently sequesters other alkaloids chemically similar to epibatidine such as N-methylepibatidine and phantasmidine (37), and it is possible that F106L protects against those. At the moment, we are unable to assign any adaptive value to the F106L substitution in *Epipedobates* nAChRs.

### Cost of S108C is Decreased Number of Receptors in the Plasma Membrane

The most parsimonious explanation for our results is that S108C causes the β2 subunit to be less efficient in its synthesis or folding, or in the assembly, trafficking and/or stability of the functional pentamer. Indeed, the β2 subunit limits the rate of receptor assembly (38, 39) and entry into the endoplasmic reticulum (40). If S108C were to cause β2 subunits to assemble poorly or assembled pentamers to be trafficked less efficiently, S108C-containing subunits would produce fewer functional receptors than an equivalent number of β2 subunits without this substitution. By the law of mass action, this would be especially evident when the number of β2 subunits is low, as we have demonstrated in this study using expression ratios of 7α:1β.

This explanation also fits the effects of S108C on mammalian α4β2 nAChRs. S108C in a mammalian β2 subunit biases the receptor toward more LS sites, probably by modifying the stoichiometry of the receptor. A β2 subunit that associates less efficiently would allow more α4 subunits to be incorporated into assembling receptors, which would result in proportionally more LS sites and a rightward shift of the ACh CRC that was evident in our previous work (Additional file 2, panels A and B).

## Conclusions

While the S108C substitution may bestow epibatidine resistance on the α4β2 nAChRs, the decrease in the numbers of receptors in the plasma membrane incurs a potential cost of decreased cholinergic synaptic transmission unless this is remedied by further adaptations. Recent evidence highlights the importance of chaperones (a protein that interacts with another protein to acquire its functionally active conformation), including nAChR-specific chaperones, in determining nAChR subunit stoichiometry and trafficking (movement of proteins within or out of the cell) in mammals (41–43). Poison frogs might compensate for S108C-dependent assembly inefficiency by regulating receptor stoichiometry or availability *via* chaperones. Other possible solutions to this problem would be up-regulation of genes or proteins for nAChR subunits or chaperones in the brain, which could be examined with transcriptomic and proteomic studies of dendrobatid brains. For example, a simple prediction from our results is that to compensate for reduced levels of α4β2 nAChRs, dendrobatids with the β2 S108C substitution must constantly generate extremely high levels of mRNA and/or protein for β2 subunits compared with dendrobatids without S108C or non-dendrobatids, to ensure an abundant population of β2-containing receptors; this would levy a continuing metabolic cost.

## Methods

### Chemicals

All chemicals and solvents were of analytical grade, purchased from Sigma-Aldrich (St. Louis, MO) and Life Technologies (Grand Island, NY). The iodinated monoclonal antibody ^125^I-mAb295 was kindly provided by Dr. Jon Lindstrom (University of Pennsylvania; Philadelphia, PA) and Dr. Paul Whiteaker (The Barrow Neurological Institute, Phoenix, AZ).

### In vitro transcription of single subunits

For experiments with single (non-concatenated) subunits, we used DNA encoding α4 and β2 nAChR subunits (chrna4 and chrnb2) from different species. DNA encoding *Xenopus* subunits cloned in pCMV-SPORT6 were obtained from Dharmacon (Lafayette, CO). DNAs encoding *Epipedobates* and *Nanorana* nAChR subunits were optimized for expression in *Xenopus laevis* oocytes, synthesized *de novo* and subcloned in pGEMHE by GenScript (Piscataway, NJ). Complementary DNAs encoding human nAChR subunits were cloned in pSP64. After linearizing the plasmids, nAChR subunits were *in vitro* transcribed using mMessage mMachine (Life Technologies, Grand Island, NY). RNA concentration and quality were checked using a ND-1000 spectrophotometer (NanoDrop Technologies, Wilmington, DE), electrophoresis (either Bioanalyzer or TapeStation systems, Agilent, Santa Clara, CA), or fluorometry (Qubit, ThermoFisher Scientific, Waltham, MA). RNA stocks were stored at −80°C and aliquots were stored at −20°C.

### Oocyte isolation and injection of single subunits

Mature *Xenopus laevis* frogs were obtained from Nasco (Fort Atkinson, WI) and housed in the University of Texas animal facility. Frogs underwent partial oophorectomy under tricaine anesthesia, and the piece of ovary was placed in isolation media (108 mM NaCl, 2 mM KCl, 1 mM EDTA, 10 mM HEPES, pH = 7.5). Using forceps, the thecal and epithelial layers were manually removed from stage V and VI oocytes. Isolated oocytes were treated with collagenase from *Clostridium histolytic* (83 mM NaCl, 2 mM KCl, 1 mM MgCl_2_, 5 mM HEPES, and 0.5 mg mL^-1^ collagenase) for 10 min to remove the follicular layer. Each oocyte was injected in the cytoplasm using a microinjector (Drummond Scientific Company, Broomall, PA) with RNA encoding nAChR single subunits in a volume of 50 nL. The subunits were either wild-type or with substitutions in either position 106 and or 108 of the β2 subunit (numeration corresponds to the mature protein). For *Xenopus* subunits, 1α4:3β2 W/W ratio: 6 ng total RNA for FS (F106, S108; wild-type) receptors and 20 ng total RNA for F**C** and **LC** mutants (the residue in bold font is the substitution); 7α4:1β2 W/W ratio: 23 ng total RNA for FS receptors and 32 ng total RNA for FC and LC receptors. For *Nanorana* subunits, 1α4:3β2 W/W ratio: 10 ng total RNA for all genotypes; 7α4:1β2 W/W ratio: 24 ng total RNA for all genotypes. For *Epipedobates* subunits, 16 ng total RNA for all genotypes and all ratios (α4:β2 in 1:7 or 7:1 W/W). For human subunits, 10 ng total RNA for all genotypes and all ratios (α4:β2 in 1:3 or 3:1 W/W). For *Xenopus* and *Nanorana*, we aimed to maximize the current and injected variable amounts of RNA depending on the construct and α:β ratio. Oocytes were then incubated at 16°C in sterile incubation solution (88 mM NaCl, 1mM KCl, 2.4 mM NaHCO_3_, 19 mM HEPES, 0.82 mM MgSO_4_, 0.33 mM Ca(NO_3_)_2_, 0.91 mM CaCl_2_, 10,000 units/L penicillin, 50 mg/L gentamicin, 90 mg/L theophylline, and 220 mg/L sodium pyruvate, pH=7.5). Incubation periods varied from three to five days.

### Electrophysiological recordings of single subunits

Responses to acetylcholine (ACh) were studied 3-5 days after injection through two-electrode voltage-clamp (Oocyte Clamp OC-725C, Warner Instruments, Hamden, CT) and digitized using a PowerLab 4/30 system (ADInstruments, Colorado Springs, CO). The oocytes were placed in a rectangular chamber and continuously perfused at a rate for 2 mL min^-1^ with Ba-ND96 + Atropine buffer (96 mM NaCl, 2 mM KCl, 1 mM BaCl_2_, 1 mM MgCl_2_, 10 mM HEPES, 1 μM atropine) at room temperature. Oocytes were clamped at −70 mV using two glass electrodes filled with 3 M KCl. All drugs were applied by bath-perfusion, and solutions were prepared the day of the application.

Concentration-response curves (CRCs) for ACh were obtained by applying increasing concentrations (20-30 s applications) with 5-15 min washout times. All responses were normalized to the maximal ACh response seen in that oocyte by assigning it a 100% value. Maximal current values (I_max_) were obtained from the CRCs or from applying a single ACh concentration determined to produced maximal current based on previous CRCs (13).

### Preparation of high-sensitivity, concatenated α4β2-nAChR DNA constructs containing human or mutant β2-nAChR subunits

The engineering and design of human α4β2-nAChRs adhered to methods previously described for concatenated nAChR DNA constructs (31, 44). Briefly, all but the first β2 subunit were absent their start codons and signal peptides, and all but the last were devoid of a stop codon. Each nAChR subunit was linked to its neighbor by a short stretch of nucleotides encoding a series of 6 or 9 (Ala-Gly-Ser)_n_ repeats, engineered to ensure a total linker length (including the C-terminal tail of the preceding subunit) of 40 ± 2 amino acids. GeneArt custom gene synthesis (Invitrogen; Waltham, MA) was used to design, synthesize, and sequence-verify optimized human (FS; control) and mutant β2 nAChR subunits (either F106L, S108C, or both). A unique set of six restriction sites either flanking the entire concatemer or approximately bisecting each linker between subunits was introduced along the concatenated sequence (31). This permitted replacement of individual subunits using standard restriction digestion and ligation methods. As previously demonstrated, the initial β2-α4 subunit pair of the α4β2-nAChR will assemble to form an orthosteric binding site between the complementary (-) face of the initial β2 subunit and the principal (+) face of the following α4 subunit (31, 45). Each α4β2-nAChR construct was subcloned into the pSGEM oocyte high expression vector and assembly of each construct was verified by restriction digest. Subunits for α4β2-nAChRs were designed in the order β2-α4-β2-α4-β2 for the human α4β2-nAChR construct (control). Accordingly, the assembled plesiomorphic human α4β2-nAChR genotype hosts orthosteric agonist binding pockets at the α4(+)/(-)β2 interfaces between the first and second and the third and fourth subunits (45).

To investigate the contributions of poison frog amino acid substitutions in a human genetic background, concatenated human α4β2-nAChR constructs were engineered with single mutant β2-nAChR subunit genotypes containing the amino acid patterns F**C** (i.e. substitution S108C) and **LC** (i.e. double substitution F106L,S108C), as identified in *Epipedobates* (13). Initially, human α4β2-nAChR construct sequences were expressed with the mutant F**C** (i.e. S108C), a sequence known to confer epibatidine resistance in frogs (13). Human α4β2-nAChR constructs were engineered with this amino acid substitution using the following stoichiometries: β2(F**C**)-α4-β2-α4-β2, β2(F**C**)-α4-β2(F**C**)-α4-β2, and β2(F**C**)-α4-β2(F**C**)-α4-β2(F**C**). Additional human α4β2-nAChR constructs were engineered to express the double substitution within the β2-nAChR subunit at three different positions: β2(**LC**)-α4-β2-α4-β2, β2(**LC**)-α4-β2(**LC**)-α4-β2, and β2(**LC**)-α4-β2(**LC**)-α4-β2(**LC**).

### Concatemer RNA preparation for oocyte injection

All concatenated α4β2-nAChR DNA plasmids were linearized with SwaI (2 hr at 37°C), treated with proteinase K (30 min at 50°C), and purified using Qiagen’s PCR Clean-up Kit (Valencia, CA), cRNAs were transcribed using the mMessage mMachine T7 kit (Applied Biosystems/Ambion, Austin, TX), and were purified using the Qiagen RNeasy Clean-up kit and stored at −80°C. RNA length and quality were confirmed on a 1% agarose gel.

### Oocyte preparation and injection of RNA encoding human concatenated subunits

For expression of the human concatenated nAChRs, *Xenopus laevis* oocytes were isolated and processed for receptor expression as described in accordance with Lucero et al. (31) and with the following modification(s): *X. laevis* oocytes were purchased from Ecocyte LLC (Austin, TX), maintained at 16°C and injected with 80 nL containing 20 ng of RNA.

### Two-electrode voltage-clamp (TEVC) recordings of human concatenated α4β2-nAChRs

Detailed methodology for obtaining ACh CRCs and I_max_ from human concatenated nAChR-injected oocytes can be found in (31, 34). Briefly, seven days post-injection (for maximal expression of concatenated DNA constructs) oocytes were voltage-clamped at −70 mV with an Axoclamp 900A amplifier (Molecular Devices, Sunnyvale, CA). Recordings were sampled at 10 kHz (low pass Bessel filter, 40 Hz; high pass filter, direct current), and the traces were extracted and analyzed using Clampfit software (Molecular Devices). Oocytes with leak currents >50 nA were discarded and not used for analysis. Drugs were applied at a flow rate of 4 mL min^-1^ using a 16-channel, gravity-fed perfusion system with automated valve control (AutoMate Scientific, Inc., Berkeley, CA). Oocytes were recorded in OR2 buffer (82.5 mM NaCl, 2.5 mM KCl, 1 mM MgCl_2_, 5 mM HEPES, pH 7.6; 22 °C) supplemented with atropine sulfate (1.5 μM) to block endogenous muscarinic responses. ACh was acutely perfused for 1 sec with a 60 sec washout between ACh applications. All CRC recording sessions included oocytes injected mutant and wildtype controls (seven oocytes per group) to account for day-to-day and batch-to-batch variability in functional expression levels.

### ^125^I-mAb 295 cell-surface labeling of α4β2-nAChRs

Surface expression levels of *Epipedobates* and human concatenated α4β2-nAChRs were quantified with ^125^I-mAb 295 in an oocyte binding assay. For consistency, wildtype and mutant α4β2-nAChRs were tested on the same day. The iodinated monoclonal antibody, ^125^I-mAb 295, specifically recognizes correctly folded human, bovine, and rodent β2-nAChR subunits (33, 46, 47) and immunolabeling protocols using this antibody have previously been described (48, 49). We determined ^125^I-mAb 295 specificity to frog α4β2-nAChR isoforms by comparing specific binding of oocytes expressing *Epipedobates* α4β2-nAChR channels against uninjected oocytes (nonspecific controls; see below). Oocytes were injected with α4:β2 unbiased RNA ratios. Following injection, oocytes were incubated for seven days prior to measuring cell-surface expression (see below). Oocytes expressing *Epipedobates* α4β2-nAChRs showed significantly higher levels of cell-surface binding compared to uninjected oocytes (Additional file 4), demonstrating that I-mAb295 recognizes *Epipedobates* β2-nAChR subunit when heterologously expressed in *Xenopus* oocytes.

For immunolabeling experiments with *Epipedobates* α4β2-nAChRs, oocytes were isolated, decollagenased, and injected as described above for single subunits. Half of the oocytes were maintained at 16°C in Austin for maximal current experiments conducted on day four after injection for comparison with other frog channels (peak current elicited with a maximal concentration of ACh, 100 μM, 20 sec, 2 mL min^-1^). The other half were shipped overnight to the Barrow Neurological Institute (Phoenix, AZ) to measure the amount of α4β2-nAChR cellsurface expression and conduct maximal current experiments (concentration: 1 mM ACh, application duration: 1 sec, flow rate: 4 mL min^-1^), both on day seven after injection. Oocytes were shipped in vials of incubation solution packed in a styrofoam box with a cold pack to keep the oocytes cool and then stored at 16°C upon arrival. Despite the travel and slight variations in methodology the observed pattern of I_max_ remained consistent.

After I_max_ experiments on day seven, oocytes were sorted into sets of seven (each expressing wildtype or mutant concatenated α4β2-nAChR isoforms) on a 24-well plate (one set per well). The accompanying OR2 buffer was aspirated from each well and replaced with 2 nM ^125^I-mAb 295 in OR2, supplemented with 10% heat-inactivated fetal bovine serum (to reduce nonspecific binding) and incubated with gentle agitation for 3 hr at 22°C. Washes were performed by aspiration of the radiolabeled solution and replacement with ice-cold OR2 supplemented with 10% heat-inactivated fetal bovine serum (2 mL well^-1^). The oocytes were then transferred to a fresh 24-well plate with the minimum possible volume of diluted radioactive solution. This wash protocol was repeated three times before transferring the oocytes to a fresh 24-well plate. Oocytes were then lysed overnight in 0.1% SDS, 0.01 N NaOH (0.5 mL), prior to scintillation counting at 85% efficiency using a Packard TriCarb 1900 Liquid Scintillation Analyzer (PerkinElmer Life Sciences; Waltham, MA). One or more wells of non-injected oocyte controls were included per assay plate to determine nonspecific binding. Nonspecific binding was subtracted from total binding determined in each of the other wells of the same plate to calculate specific binding. Surface expression for human concatenated mutant channels were determined similarly, modified with preparation as described for concatenated channels and no tests of I_max_ on day four.

### Statistical analysis

Results were expressed as mean ± SEM. All statistical analysis was performed with Prism 8 (GraphPad Software Inc., San Diego, CA). For CRCs, the mean percent of maximal current for each concentration were plotted and fitted to monophasic (Eq. 1) or biphasic (Eq. 2) logistic equations. The best fitted equation was determined by applying the Extra sum-of-squares F test. The corresponding F(DFn,DFd) values corresponding to each comparison are listed in Additional 9; a significance level of p < 0.01 was used so that the analysis could properly differentiate between the two models. Once the correct equation was identified, all parameters (log EC_50_, Hill coefficients, and maximal currents) were determined. All relevant CRC parameters are shown in Additional files 6-9.

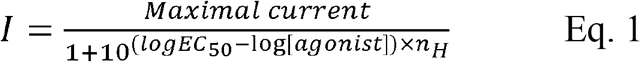

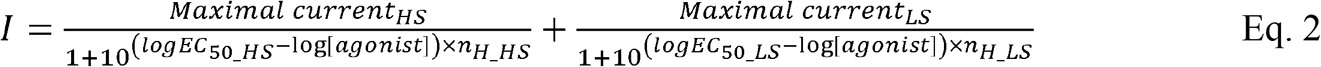

*I*, measured current; *Maximal current* achieved, expressed as a percentage of the maximal ACh response recorded in that oocyte; *EC_50_*, effective concentration 50, or concentration that produces half the maximal current; *n_H_*, Hill coefficient. In the case of biphasic curves (which reflect two kinds of binding sites), subscripts HS and LS were used to identify the two kinds of populations (HS, high sensitivity; LS, low sensitivity).

In the ^125^I-mAb 295 immunolabeling experiments, total cell-surface binding was determined for each individual experiment (seven pooled oocytes/experimental group/experimental day) then averaged across three experimental days. For each experiment, uninjected oocytes were used to calculate non-specific binding. Specific counts were determined by subtracting the average non-specific counts from the average total binding (31, 34).

Maximal currents and specific binding were analyzed using One or Two-way analysis of variance (ANOVA), followed by pairwise comparisons corrected for multiple comparisons as indicated in the legend. A significance level of p< 0.05 was used for these analyses.

## Abbreviations

ACh: acetylcholine;
β2(FC): phenylalanine in position 106 and cysteine in position 108 in the β2 subunit;
β2(FS): phenylalanine in position 106 and serine in position 108 in the β2 subunit;
β2(LC): leucine in position 106 and cysteine in position 108 in the β2 subunit;
β2(LS): leucine in position 106 and serine in position 108 in the β2 subunit;
HS: high sensitivity;
LS: low sensitivity;
mya: million years ago;
nAChR: nicotinic acetylcholine receptor;

## Additional files

### Additional file 1.pdf

Subunit arrangements and structure of α4β2 nAChRs in different stoichiometries.

(A and B). Diagrams of nAChRs in different stoichiometries, seen from the extracellular side. (A) nAChR formed by two *α4* and three β2 subunits (2α:3β) that possesses high sensitivity (HS) binding sites for ACh located at α(+):β(-) interfaces. (B) nAChR formed by three α4 and two β2 subunits (3α:2β). In addition to the HS binding sites, it possesses a low sensitivity (LS) binding site for ACh located at the interface between two adjacent α subunits, α(+):α(-). **+** and **–** signs indicate the principal and complementary components of the subunit interfaces. (C-E). Structure of human α4β2 nAChRs determined by cryo-electron microscopy (1) with bound ligand and antibody fragments. (C) Stoichiometry 2α:3β (Protein Data Bank, PDB: 6CNJ), viewed from the extracellular side. (D and E) Stoichiometry 3α:2β (PDB: 6CNK), viewed from the membrane side (D) and the extracellular side (E). Molecular graphics performed with UCSF Chimera (2). Alpha subunit in salmon, β subunit in grey, Fragment antigen-binding (Fab) from monoclonal antibodies in cyan. The arrows indicate the interfaces where nicotine (black, present in the structure) and acetylcholine bind. Red arrows indicate HS binding sites and blue arrows indicate LS binding site.

1. Walsh RM, Jr., Roh SH, Gharpure A, Morales-Perez CL, Teng J, Hibbs RE. Structural principles of distinct assemblies of the human alpha4beta2 nicotinic receptor. Nature. 2018;557(7704):261-5.
2. Pettersen EF, Goddard TD, Huang CC, Couch GS, Greenblatt DM, Meng EC, et al. UCSF Chimera--a visualization system for exploratory research and analysis. J Comput Chem. 2004;25(13):1605-12.

### Additional file 2.pdf

Effects of reciprocal substitutions on ACh concentration-response curves (CRC) in the β2 subunit of human and dendrobatid frog, *Epipedobates anthonyi*.

Data redrawn from (1), presented as mean ± SD. (A) A high ratio of β2 to α4 (1α:3β) of cRNA of the wild type human receptor subunits produces a monophasic CRC with a single EC_50_ indicating only high sensitivity (HS) binding sites (black curve; β2(FS) represents F106 and S108 in β2 subunit). Introduction of the S108C substitution adds a low sensitivity (LS) binding site so that the CRC is now best fit with a biphasic curve reflecting both HS and LS sites (green curve: β2(F**C**), amino acid in bold indicates a substitution). Further addition of F106L to S108C eliminates the LS sites, thus compensating for the effect of S108C [orange curve: β2(**LC**)]. (B) A low ratio of β2 to α4 (3α:1β) of the wild type human receptor subunits produces an ACh CRC shifted rightward and best fit with a monophasic curve with a shallow slope [black curve: β2(FS)]. Introduction of the S108C substitution shifted the curve further right (green curve: β2(F**C**)]. Addition of F106L to S108C partially compensates for the effect of S108C alone [orange curve: β2(**LC**)]. (C) When the ratio of injected human β subunit cRNA/α subunit cRNA is high, the nAChR stoichiometry is 2α:3β. However, with paucity of β subunits the stoichiometry shifts to 3α:2β. (D,E) Even with more extreme ratios of α and β subunits (1:7 and 7:1) and the introduction of the ancestral amino acids (**FS**, **F**C), there was no change in the CRC of *Epipedobates anthonyi*. (F) The stoichiometry of frog nAChR receptors is unknown but we conjectured it is 2α:3β because they show a single kind of binding site (HS). This conjecture is noted by the question mark over the grey arrow.

1. Tarvin RD, Borghese CM, Sachs W, Santos JC, Lu Y, O’Connell LA, et al. Interacting amino acid replacements allow poison frogs to evolve epibatidine resistance. Science. 2017;357(6357):1261-6.

### Additional file 3.pdf

Maximal currents from α4β2 nAChR of two species of non-dendrobatids

(A) *Xenopus tropicalis* (n= 10-39) (B) *Nanorana parkeri* (n= 9-20). The number over each bar indicates the total amount of cRNA (ng) injected per oocyte, while maintaining the α:β RNA ratio indicated. β2(FS) represents F106 and S108 in the β2 subunit. β2(F**C**) and β2(**LC**) indicates the residues present in position 106 and 108 in the β2 subunit, with the bold font indicating substitutions in the wild type background.

### Additional file 4.pdf

Antibody verification. Oocytes were injected with cRNA encoding *Epipedobates anthonyi* α4β2 nAChRs (ratio 1:1, 4 ng each).

A) Currents induced by 1 mM ACh 7 days after injection (n=21); uninjected oocytes were assumed to have no response to ACh based on previous experiments. B) Raw counts obtained with an iodinated antibody directed against the β2 subunit (^125^I-mAb 295) in each group (n=3 experiments with 7 pooled oocytes per experiment). C) Correlation between the maximal ACh-induced current and the specific binding observed for each of the pooled oocytes expressing *Epipedobates* nAChRs tested for this study (R^2^= 0.83). β2(LC) represents L106 and C108 in the β2 subunit. When used for residues, the bold font indicates substitutions in the wild type background.

Uninjected oocytes were used as blanks, and their counts subtracted from the values of injected oocytes for each experiment.

### Additional File 5.pdf

Accession numbers and species names included in Fig 1.

The names of undefended species of poison frogs (Dendrobatidae) are in black and those of defended species are in blue.

### Additional File 6.pdf

Parameters from non-linear curve fit of ACh concentration-response curves in oocytes expressing *Xenopus tropicalis* α4β2 nAChRs.

The data from all curves fitted a one-population curve. β2(FS) represents F106 and S108 in the β2 subunit. When used for residues, the bold font indicates substitutions in the wild type background.

### Additional File 7.pdf

Parameters from non-linear curve fit of ACh concentration-response curves in oocytes expressing *Nanorana* α4β2 nAChRs.

The data was best fitted by a biphasic curve. β2(FS) represents F106 and S108 in the β2 subunit.

When used for residues, the bold font indicates substitutions in the wild type background.

### Additional File 8.pdf

Parameters from non-linear curve fit of ACh concentration-response curves in oocytes expressing concatemeric human α4β2 nAChRs (2 α4:3 β2).

The data from each curve fit a monophasic curve. β2(F**C**) represents F106 and C108 in the β2 subunit. The bold font indicates a substitution in the wild type background. P1, P3, P5 refer to the position of the β2 subunit in the concatemer (Fig 4).

### Additional File 9.pdf

F(DFn, DFd) and p values from one- and two population fittings to ACh concentration-response curves, calculated using the Extra sum-of-squares F test.

A value of *P* < 0.10 means the preferred model is the “Two populations” (biphasic curve); *P*> 0.10 means the preferred model is the “One population” (monophasic curve). DFn, degree of freedom for the numerator of the F ratio, DFd is for the denominator. The amino acids between parentheses stand for the residues at locations 106 and 108, respectively; if there is a substitution, the letter is in bold font.

## Declarations

### Ethics approval and consent to participate

Frog care and surgery were conducted under IACUC approved protocols (AUP-2015-00205 and AUP-2016-00016).

### Consent for publication

Not applicable

### Availability of data and materials

The datasets used and/or analyzed during the current study are available from the corresponding author on reasonable request.

### Author Contributions

Conceptualization: H.H.Z., D.C.C., J.M.Y., C.M.B., and A.A.G. Project administration: H. H.Z. Investigation: J.M.Y. and A.A.G. Formal analysis: J.M.Y., C.M.B., A.A.G. Visualization: C.M.B., A.A.G., D.C.C. Writing – Original Draft Preparation: H.H.Z., C.M.B., A.A.G, J.M.Y. Writing – Review & Editing: H.H.Z., D.C.C., A.A.G., C.M.B., J.M.Y. Funding acquisition: H.H.Z., D.C.C., A.A.G., J.M.Y.

## Acknowledgments

We thank Drs. Jon Lindstrom for his generous gift of iodinated antibody, John Mihic for the use of his lab space and equipment, and Sonia Mason and Jelena Todorovic for experimental assistance.

