## Additional file 1 for "A potential cost of evolving epibatidine resistance in poison frogs"

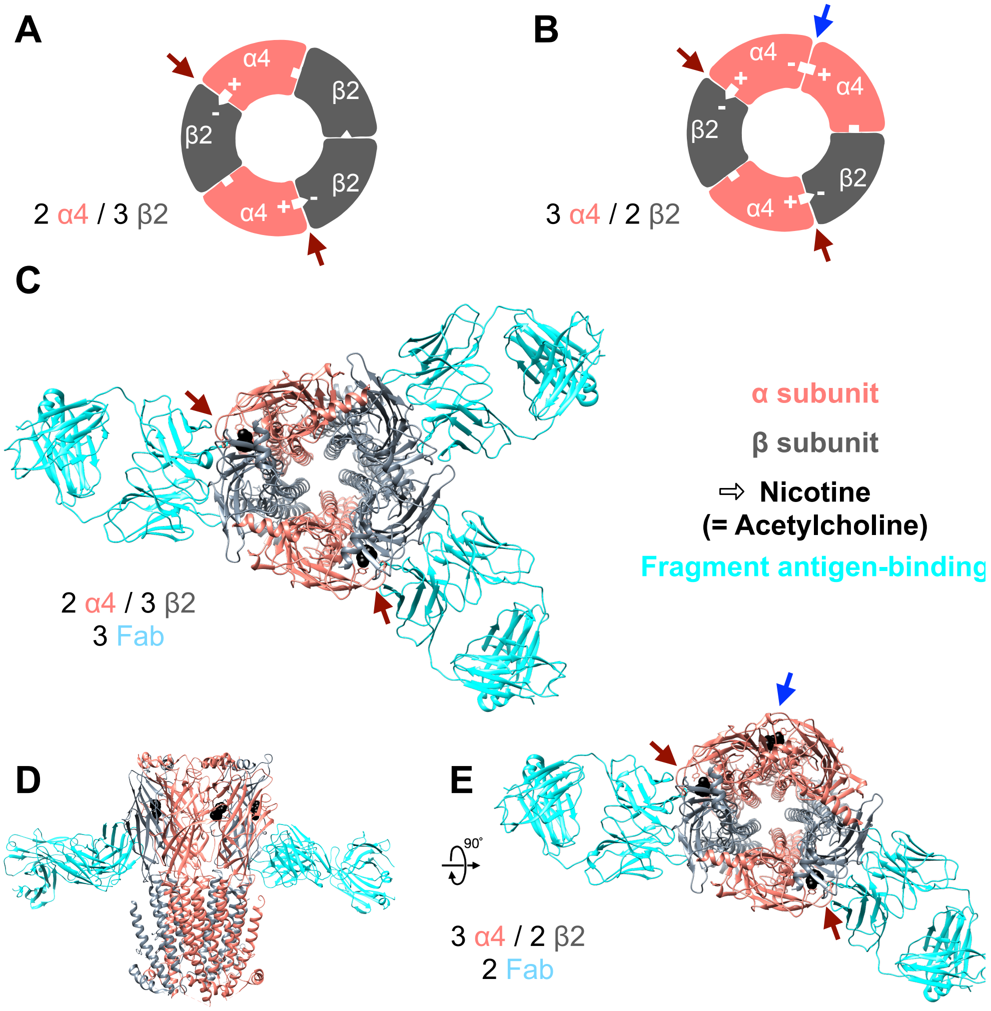


**Additional File 1. Subunit arrangements and structure of α4β2 nAChRs in different stoichiometries.** (A and B). Diagrams of nAChRs in different stoichiometries, seen from the extracellular side. (A) nAChR formed by two α4 and three β2 subunits (2α:3β) that possesses high sensitivity (HS) binding sites for ACh located at α(+):β(−) interfaces. (B) nAChR formed by three α4 and two β2 subunits (3α:2β). In addition to the HS binding sites, it possesses a low sensitivity (LS) binding site for ACh located at the interface between two adjacent α subunits, α(+):α(-). + and – signs indicate the principal and complementary components of the subunit interfaces. (C-E). Structure of human α4β2 nAChRs determined by cryo-electron microscopy (1) with bound ligand and antibody fragments. (C) Stoichiometry 2α:3β (Protein Data Bank, PDB: 6CNJ), viewed from the extracellular side. (D and E) Stoichiometry 3α:2β (PDB: 6CNK), viewed from the membrane side (D) and the extracellular side (E). Molecular graphics performed with UCSF Chimera (2). Alpha subunit in salmon, β subunit in grey, Fragment antigen-binding (Fab) from monoclonal antibodies in cyan. The arrows indicate the interfaces where nicotine (black, present in the structure) and acetylcholine bind. Red arrows indicate HS binding sites and blue arrows indicate LS binding site.

1. Walsh RM, Jr., Roh SH, Gharpure A, Morales-Perez CL, Teng J, Hibbs RE. Structural principles of distinct assemblies of the human alpha4beta2 nicotinic receptor. Nature. 2018;557(7704):261-5.
2. Pettersen EF, Goddard TD, Huang CC, Couch GS, Greenblatt DM, Meng EC, et al. UCSF Chimera--a visualization system for exploratory research and analysis. J Comput Chem. 2004;25(13):1605-12.
