## Additional file 2 for "A potential cost of evolving epibatidine resistance in poison frogs"

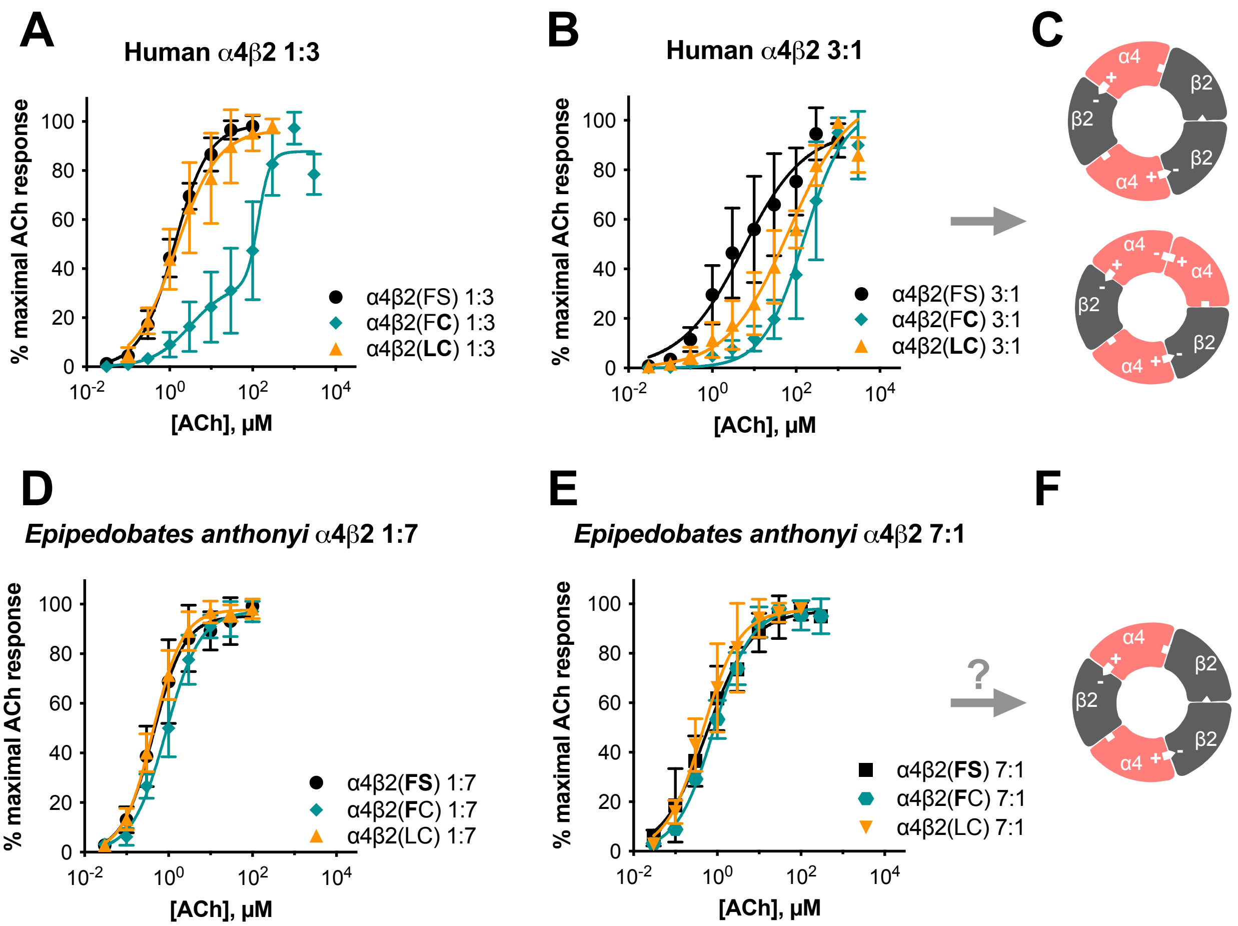


**Additional File 2. Effects of reciprocal substitutions on ACh concentration-response curves (CRC) in the β2 subunit of human and dendrobatid frog, *Epipedobates anthonyi*.**

Data redrawn from (1), presented as mean ± SD. (A) A high ratio of β2 to α4 (1α:3β) of cRNA of the wild type human receptor subunits produces a monophasic CRC with a single $EC_{50}$ indicating only high sensitivity (HS) binding sites (black curve; β2(FS) represents F106 and S108 in β2 subunit). Introduction of the S108C substitution adds a low sensitivity (LS) binding site so that the CRC is now best fit with a biphasic curve reflecting both HS and LS sites (green curve: β2(F**C**), amino acid in bold indicates a substitution). Further addition of F106L to S108C eliminates the LS sites, thus compensating for the effect of S108C [orange curve: β2(**LC**)]. (B) A low ratio of β2 to α4 (3α:1β) of the wild type human receptor subunits produces an ACh CRC shifted rightward and best fit with a monophasic curve with a shallow slope [black curve: β2(FS)]. Introduction of the S108C substitution shifted the curve further right (green curve: β2(F**C**)]. Addition of F106L to S108C partially compensates for the effect of S108C alone [orange curve: β2(**LC**)]. (C) When the ratio of injected human β subunit cRNA/α subunit cRNA is high, the nAChR stoichiometry is 2α:3β. However, with paucity of β subunits the stoichiometry shifts to 3α:2β. (D,E) Even with more extreme ratios of α and β subunits (1:7 and 7:1) and the introduction of the ancestral amino acids (**FS**, **FC**), there was no change in the CRC of *Epipedobates anthonyi*. (F) The stoichiometry of frog nAChR receptors is unknown but we conjectured it is 2α:3β because they show a single kind of binding site (HS). This conjecture is noted by the question mark over the grey arrow.


1) Tarvin RD, Borghese CM, Sachs W, Santos JC, Lu Y, O'Connell LA, et al. Interacting amino acid replacements allow poison frogs to evolve epibatidine resistance. Science. 2017;357(6357):1261-6.
