## Additional file 3 for "A potential cost of evolving epibatidine resistance in poison frogs"

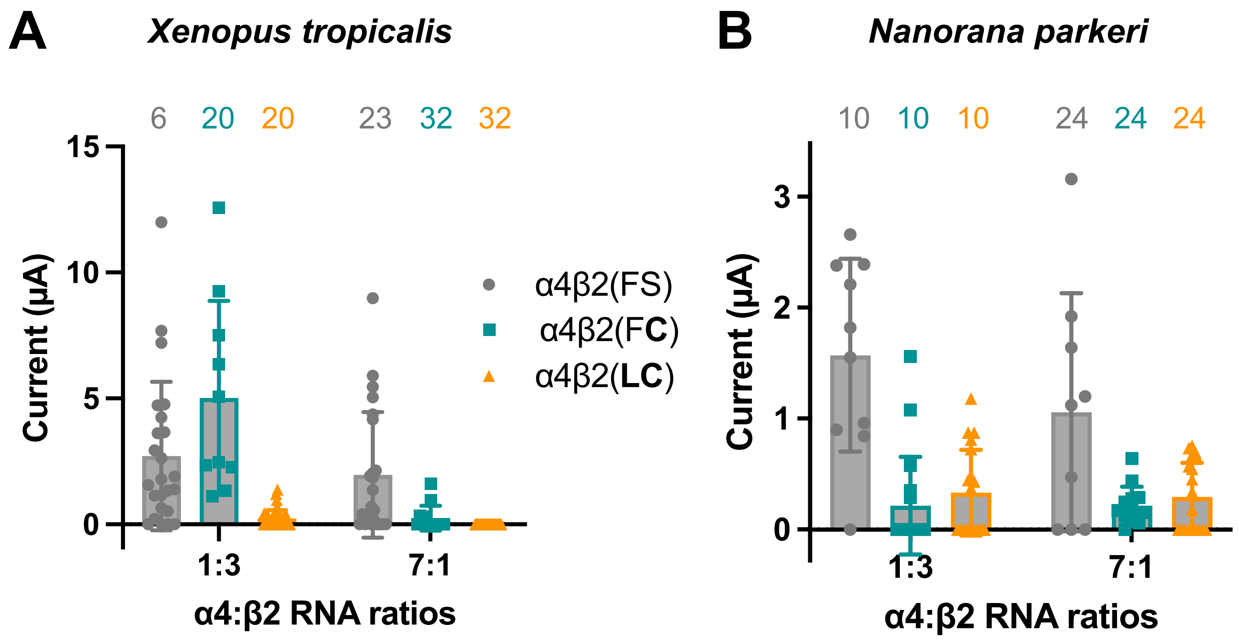


**Additional File 3. Maximal currents from α4β2 nAChR of two species of non-dendrobatids**. (A) *Xenopus tropicalis* (n= 10-39) (B) *Nanorana parkeri* (n= 9-20). The number over each bar indicates the total amount of cRNA (ng) injected per oocyte, while maintaining the α:β RNA ratio indicated. β2(FS) represents F106 and S108 in the β2 subunit. β2(F**C**) and β2(**LC**) indicates the residues present in position 106 and 108 in the β2 subunit, with the bold font indicating substitutions in the wild type background.
