## Additional file 4 for "A potential cost of evolving epibatidine resistance in poison frogs"

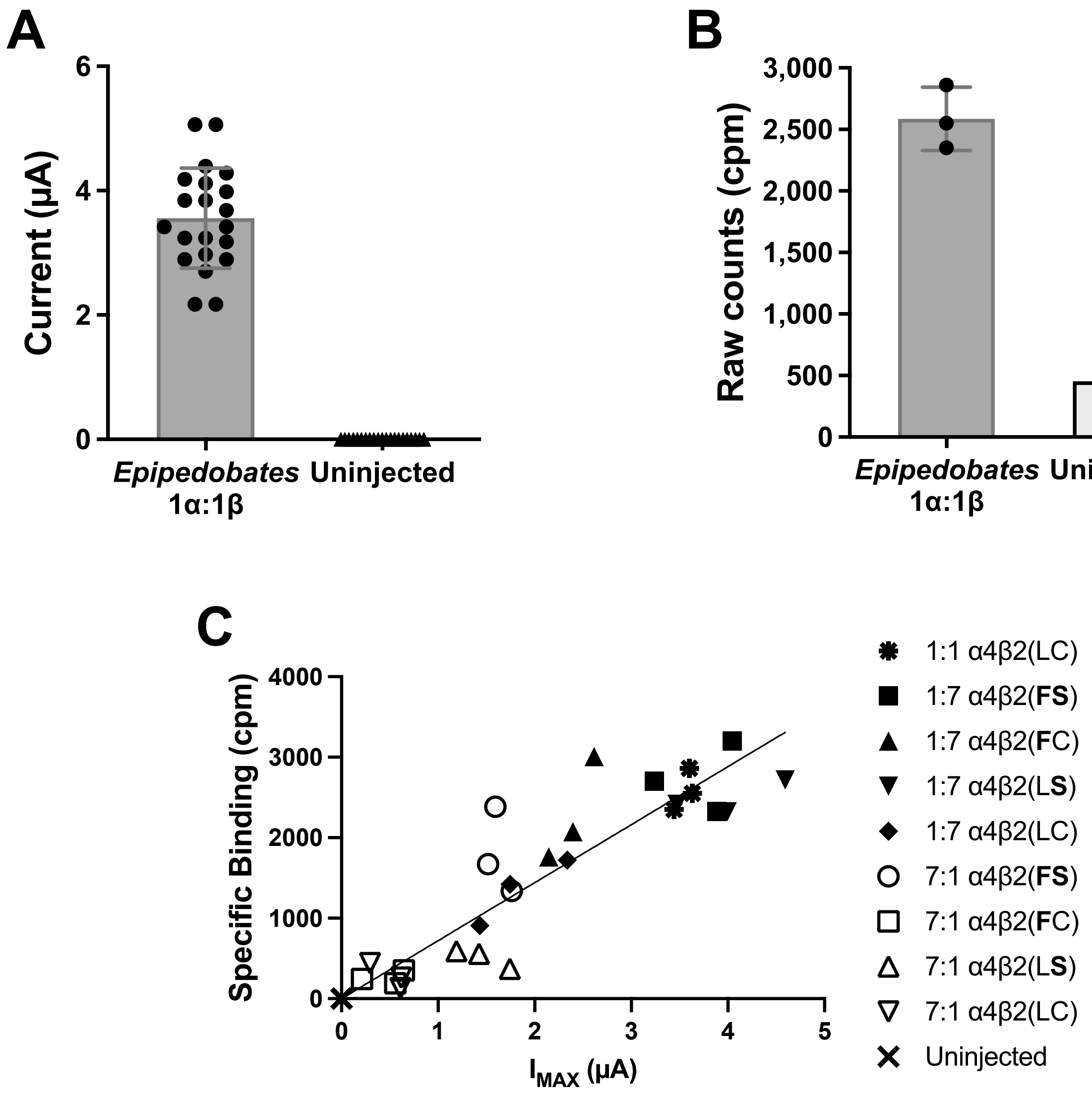


**Additional File 4.** Antibody verification. Oocytes were injected with cRNA encoding *Epipedobates anthonyi* α4β2 nAChRs (ratio 1:1, 4 ng each).
A) Currents induced by 1 mM ACh 7 days after injection (n=21); uninjected oocytes were assumed to have no response to ACh based on previous experiments. B) Raw counts obtained with an iodinated antibody directed against the β2 subunit ($^{125}$I-mAb 295) in each group (n=3 experiments with 7 pooled oocytes per experiment). C) Correlation between the maximal ACh-induced current and the specific binding observed for each of the pooled oocytes expressing *Epipedobates* nAChRs tested for this study ($R^2$= 0.83). β2(LC) represents L106 and C108 in the β2 subunit. When used for residues, the bold font indicates substitutions in the wild type background.
Uninjected oocytes were used as blanks, and their counts subtracted from the values of injected oocytes for each experiment.
