## Additional file 5 for "A potential cost of evolving epibatidine resistance in poison frogs"

**Additional File 5**. **Accession numbers and species names included in Fig 1.** The names of undefended species of poison frogs (Dendrobatidae) are in black and those of defended species are in blue.

| Subunit | Common name | Species | Accession number |
|---|---|---|---|
| β2 | tunicate | *Ciona intestinalis* | NP_001265876.1 |
| | sea lamprey | *Petromyzon marinus* | XP_032806233.1 |
| | great white shark | *Charcharodon carcharias* | XP_041035646.1 |
| | zebrafish | *Danio rerio* | XP_005169811.1 |
| | human | *Homo sapiens* | CAD88996.1 |
| | chicken | *Gallus gallus* | NP_990144.1 |
| | Western clawed frog | *Xenopus tropicalis* | NP_001093684.1 |
| | High Himalaya frog | *Nanorana parkeri* | XP_018425584.1 |
| | grass frog | *Rana temporaria* | XP_040188232.1 |
| | brilliant-thighed poison frog | *Allobates femoralis* | ATG31806.1 |
| | | *Hyloxalus italoi* | ATG31785.1 |
| | strawberry poison frog | *Oophaga pumilio* | ATG71844.1 |
| | Rio Santiago poison frog | *Excidobates captivus* | ATG31798.1 |
| | golden poison frog | *Phyllobates terribilis* | ATG31810.1 |
| | Ecuador poison frog | *Ameerega bilinguis* | ATG31790.1 |
| | | *Silversotneia cf. gutturalis* | ATG31801.1 |
| | | *Leucostethus fugax* | ATG31797.1 |
| | Anthony's poison frog | *Epipedobates anthonyi* | ATG31796.1 |
| α4 | human | *Homo sapiens* | AAB40111.1 |
| | Western clawed frog | *Xenopus tropicalis* | NP_001107315.1 |
| | High Himalaya frog | *Nanorana parkeri* | XP_018415603.1 |
| | Anthony's poison frog | *Epipedobates anthonyi* | ATG31830.1 |
