## Additional file 6 for "A potential cost of evolving epibatidine resistance in poison frogs"

**Additional File 6. Parameters from non-linear curve fit of ACh concentration-response curves in oocytes expressing *Xenopus tropicalis* α4β2 nAChRs.**

The data from all curves fitted a one-population curve. β2(FS) represents F106 and S108 in the β2 subunit. When used for residues, the bold font indicates substitutions in the wild type background.

| cRNA Ratio | Receptor | $EC_{50}$ (nM) | $n_H$ | $I_{MAX}$ (%) | n |
|---|---|---|---|---|---|
| **1:3** | α4β2(FS) | 47 (37 to 61) | 0.85 ± 0.06 | 97 ± 2 | 18 |
| | α4β2(F**C**) | 18 (13 to 26) | 1.3 ± 0.2 | 92 ± 3 | 14 |
| | α4β2(**LC**) | 21 (12 to 43) | 2.2 ± 0.8 | 83 ± 4 | 6 |
| **7:1** | α4β2(FS) | 14 (8 to 30) | 1.0 ± 0.2 | 90 ± 4 | 13 |
| | α4β2(F**C**) | 20 (17 to 24) | 1.5 ± 0.1 | 94 ± 1 | 10 |
| | α4β2(**LC**)* | - | - | - | 37 |

$EC_{50}$: effective concentration 50 (concentration that produces 50% of maximal response). Expressed as mean (95% confidence intervals).

$n_H$: Hill coefficient. Expressed as mean ± SEM.

$I_{MAX}$: Maximal current, expressed as a percentage of maximal response (mean ± SEM).

n: number of oocytes.

*Currents too small to determine concentration-response curves
