## Additional file 7 for "A potential cost of evolving epibatidine resistance in poison frogs"

**Additional File 7. Parameters from non-linear curve fit of ACh concentration-response curves in oocytes expressing *Nanorana* α4β2 nAChRs.**

The data was best fitted by a biphasic curve. β2(FS) represents F106 and S108 in the β2 subunit. When used for residues, the bold font indicates substitutions in the wild type background.

| **cRNA Ratio** | **Receptor** | **$EC_{50\_HS}$ (µM)** | **$EC_{50\_LS}$ (µM)** | **$n_{H\_HS}$** | **$n_{H\_LS}$** | **$I_{MAX\ HS}$ (%)** | **$I_{MAX\ LS}$ (%)** | **n** |
|---|---|---|---|---|---|---|---|---|
| **1:3** | α4β2(FS) | 0.29 (0.18 to 0.49) | 102 (63 to 231) | 1.4 ± 0.3 | 1.4 ± 0.4 | 47.1 ± 3.3 | 54.7 ± 6.3 | 9 |
| | α4β2(F**C**) | 0.63 (0.30 to 1.40) | 935 (92 to ?) | 2.0 ± 0.7 | 0.7 ± 0.6 | 41.1 ± 9.7 | 107 ± 150 | 5 |
| | α4β2(**LC**) | 0.41 (0.32 to 0.55) | 269 (201 to 530) | 1.6 ± 0.2 | 1.2 ± 0.1 | 35.0 ± 1.4 | 79.1 ± 6.5 | 8 |
| **7:1** | α4β2(FS) | 0.43 (0.29 to 0.64) | 117 (92 to 151) | 1.5 ± 0.2 | 1.6 ± 0.2 | 40.0 ± 1.8 | 62.2 + 3.5 | 6 |
| | α4β2(F**C**)* | - | - | - | - | - | - | 16 |
| | α4β2(**LC**) | 0.33 (? to 0.35) | 184 (173 to 196) | 4.6 ± 1.3 | 1.7 ± 0.1 | 38.7 ± 0.2 | 64.8 ± 0.7 | 9 |

$EC_{50}$: effective concentration 50 (concentration that produces 50% of maximal response). Expressed as mean (95% confidence intervals).

$n_H$: Hill coefficient. Expressed as mean ± SEM.

$I_{MAX}$: Maximal current, expressed as a percentage of maximal response (mean ± SEM).

n: number of oocytes.

The two components of the biphasic curve are characterized by different parameters. HS stands for high sensitivity, and LS for low sensitivity.

*Currents too small to determine concentration-response curves
