## Additional file 8 for "A potential cost of evolving epibatidine resistance in poison frogs"

**Additional File 8. Parameters from non-linear curve fit of ACh concentration-response curves in oocytes expressing concatemeric human α4β2 nAChRs (2 α4:3 β2).**

The data from each curve fit a monophasic curve. β2(F**C**) represents F106 and C108 in the β2 subunit. The bold font indicates a substitution in the wild type background. P1, P3, P5 refer to the position of the β2 subunit in the concatemer (Fig 4).

| **Receptor** | **$EC_{50}$ (μM)** | **$n_H$** | **$I_{MAX}$ (%)** | **n** |
|---|---|---|---|---|
| α4β2 Control | 1.3 (1.2 to 1.5) | 1.04 ± 0.05 | 96 ± 1 | 14 |
| α4β2(F**C**) P3 | 1.4 (1.3 to 1.5) | 1.06 ± 0.03 | 98 ± 1 | 14 |
| α4β2(F**C**) P1,P3 | - | - | 0 | 7 |
| α4β2(F**C**) P1,P3,P5 | - | - | 0 | 7 |

$EC_{50}$: effective concentration 50 (concentration that produces 50% of maximal response). Expressed as mean (95% confidence intervals).

$n_H$: Hill coefficient. Expressed as mean ± SEM.

$I_{MAX}$: Maximal current, expressed as a percentage of maximal response (mean ± SEM).

n: number of oocytes.
